# The abundance of Nudix hydrolase 16 transcripts is elevated in human sepsis: perspectives gained during a hands-on reductionist investigation workshop

**DOI:** 10.1101/490565

**Authors:** Mathieu Garand, Darawan Rinchai, Basirudeen Syed Ahamed Kabeer, Mohammed Toufiq, Mohamed Alfaki, Jessica Roelands, Wouter Hendrickx, Sabri Boughorbel, Nico Marr, Davide Bedognetti, Damien Chaussabel

## Abstract

A hands-on training workshop was devised with omics data used as source material to emulate reductionist investigation approaches. NUDT16, a member of the Nudix hydrolase family was selected as a candidate gene on the basis of: 1) it being upregulated in neutrophils exposed in vitro to serum of patients with sepsis, AND 2) the absence of overlap between the NUDT16 and sepsis, inflammation or neutrophil literature. We next sought to corroborate the initial finding in five public transcriptome sepsis datasets in which NUDT16 transcript levels were measured. In each of these dataset NUDT16 transcript abundance was found to be significantly increased in septic patients in vivo in comparison to uninfected controls. Next, biological concepts were extracted from the NUDT16 literature. The main concepts to emerge from profiling this literature were RNA decapping, inosine triphosphate, and inosine diphosphate. Through these concepts, indirect links could be established between NUDT16 and the sepsis/inflammation/neutrophil literature. A potential role for NUDT16 could, in turn, be inferred in the degradation of mRNAs in activated neutrophils. Follow on experiments that would be necessary in order to further explore such inference are discussed.

## INTRODUCTION

The findings described in this short discovery note were made during the preparation of training material for workshops conducted by Sidra Medicine investigators in November and December of 2018. This activity is primarily designed to help trainees build skills used while conducting reductionist investigations. Available public transcriptome datasets were used to emulate steps such as candidate gene selection, assessment of knowledge gaps, validation and interpretation/hypothesis formation. A recent review provides an overview of the program this training module is part of [1]. More details regarding the format of this “COD1 workshop” are also provided in a separate communication along with relevant training material (manuscript in preparation).

An increase in abundance of the NUDT16 transcript following exposure of neutrophils to septic serum was the starting point for the training activity. NUDT16 is a member of the Nudix hydrolase family. It includes enzymes catabolizing nucleoside triphosphates, nonnucleoside polyphosphates, as well as capped mRNAs [2]. In mammalian cells the decapping activity occurs in both the nucleolus and cytoplasm. Hydrolysis of the RNA cap by NUDT16 is part of the the exonuclease degradation pathway which initiates RNA turnover [3]. The activity of NUDT16 is similar to that of the Nudix domain protein DCP2 (Decapping MRNA 2).

Sepsis is caused by a systemic inflammatory response resulting from an aberrant host response to infection [4]. Sepsis is considered severe when it leads to multiple organ dysfunction. Septic shock is used to designate cases where sepsis is accompanied by refractory hypotension. Sepsis is associated with a strong activation of the innate immune system that is mediated by the recognition of pathogen-derived molecules that engage receptors expressed by a wide range of host cells [5]. Complement activation and activation of the coagulation cascade are two additional hallmarks of the pathogenesis process that unravels during sepsis [6]. It is accompanied by the activation of vascular endothelial cells, neutrophils and platelets. Neutrophils contribute to the innate immune response that controls of the infection but also results in considerable collateral tissue damage [7]. Neutrophil activity is mediated through the release of soluble inflammatory mediators and microbicidal molecules, as well as neutrophil extracellular traps (NETs), which are composed of DNA, histones and serine proteases [8]. NETs entrap pathogens and contribute to their elimination. However, their presence in the circulation of septic patients has also been associated with thrombosis and organ dysfunction [9].

## RESULTS

### Abundance of NUDT16 transcripts is increased in neutrophils in vitro in response to septic serum

NUDT16 was selected among a set of genes which abundance was increased following exposure of neutrophils with serum of patients with sepsis (the data was generated by our group and is publically available in GEO under accession GSE49755 [10]).

In this experiment, which is described in **Figure 1A**, neutrophils were isolated from healthy donors and cultured for 6 hours in the presence of serum obtained from other healthy individuals or individuals hospitalized with bacterial sepsis. Two different neutrophil donors where used and exposed to sera of 6 healthy controls (12 samples total) and 12 septic patients (24 samples total). Two control conditions were also included: culture without serum, with or without LPS. Transcriptional profiles were generated using Illumina Beadarrays. The dataset has been uploaded into a custom data-browsing application, GXB (<u>link</u>). NUDT16 expression profile is shown in **Figure 1B**. It is also accessible via the GXB application which includes all available sample information (link). Expression data along with sample information is also available in a spreadsheet format (supplemental File 1). Significant differences were found in both levels of expression and variance between the control serum and septic serum treated groups (t-test p<0.005, F-test <0.001).

**Figure 1:**
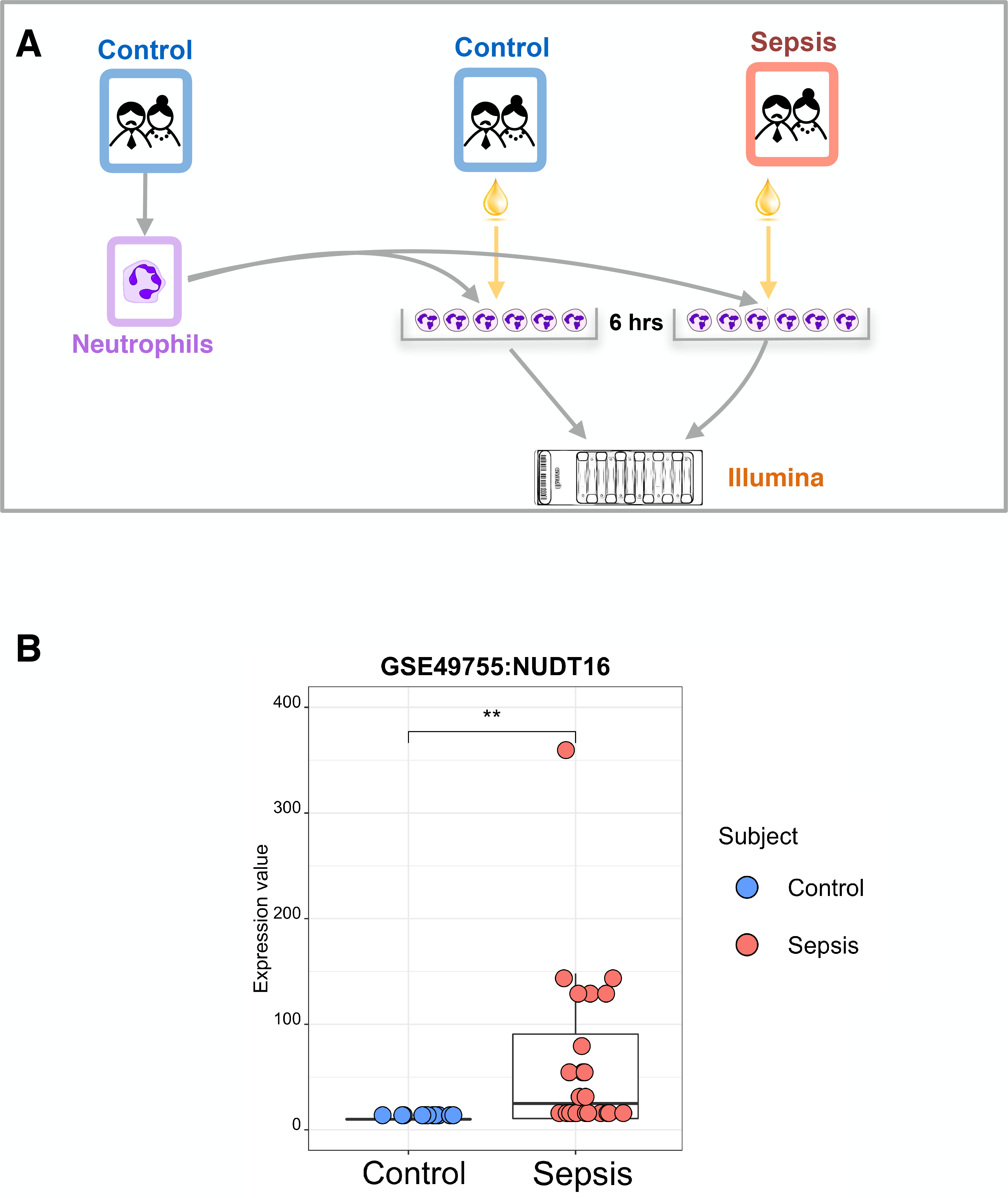
Primary observation. A. Design of the primary dataset: A dataset deposited by our group in the NCBI GEO public repository (GSE49755) was used as a starting point for selection of a gene candidate. The study was conducted in the North East of Thailand. It involved adult subjects who were hospitalized with symptoms of sepsis and showed culture positive results for either the pathogen *Burkholderia pseudomallei*, the etiological agent of melioidosis, or other bacterial species. Serum was fractionated from the blood of those patients, as well as from non-infected controls and used in in vitro cultures of neutrophils isolated from two healthy donors. After 6 hours of cultures cells were lysed and RNA extracted to be subsequently profiled on Illumina HT12 Beadarrays. B. Expression profile of the candidate gene NUDT16. Expression levels of the Nudix Hydrolase family member 16 gene after exposure in vitro of neutrophils to uninfected or septic serum is shown. Expression was measured via microarray profiling.

### No known role for NUDT16 in the context of sepsis, inflammation or neutrophil immunobiology

The next step consisted in assessing whether reports in the literature could be found that describe a role for NUDT16 in the context of sepsis (knowledge gap assessment). The NUDT16 literature was retrieved using a PubMed query which comprises its official symbol, name and known aliases:

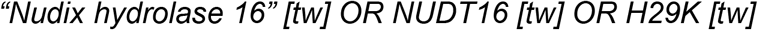

As of December of 2018 this query returned 26 results. No overlap was found with the sepsis literature (166,533 PubMed entries). Extending the search to the literature on Inflammation or Neutrophils did not return any overlapping articles either (633,921, and 116,871 PubMed entries, respectively) (**Supplementary Figure 1**).

### Increase in abundance of NUDT16 transcripts in the context of sepsis is validated in independent datasets

Next, datasets were sought in GEO that could be used to validate independently the initial finding in a relevant clinical setting. In order to do so, datasets were selected without a priori knowledge of NUDT16 expression levels. All six datasets selected were from human studies in which transcriptome profiles were generated in septic patients and compared with uninfected controls (**Table 1**). Then NUDT16 expression levels were accessed and significance levels determined based on comparison of control and sepsis cases (**Figure 2A**). NUDT16 expression was not measured by the array used in one of those six datasets. Significant increases in NUDT16 levels were found in four out of five remaining ones. Levels in abundance were not significantly different in one out of the five. Notably the fold changes that were recorded were lower than that observed in the primary dataset (ranging from 1.3 to 2.1 fold changes vs 5.7 fold changes in the septic plasma exposure dataset). This may be attributed to the fact that the validation datasets were all from in vivo studies and in mixed populations, which except for one datasets consisted of whole blood or PBMCs. Robustness of the increase in abundance of NUDT16 transcript in the context of sepsis can be further ascertain from the range of settings and methodologies employed across the validation datasets. **Figure 2B** indicates the geographic localization of the patient populations under study, which span four continents; the type of biological samples ranges from purified neutrophils, Peripheral Blood Mononuclear Cells (PBMCs) and whole blood. Studies include both pediatric and adult populations. Finally, data were generated using two different microarray platforms.

**Figure 2:**
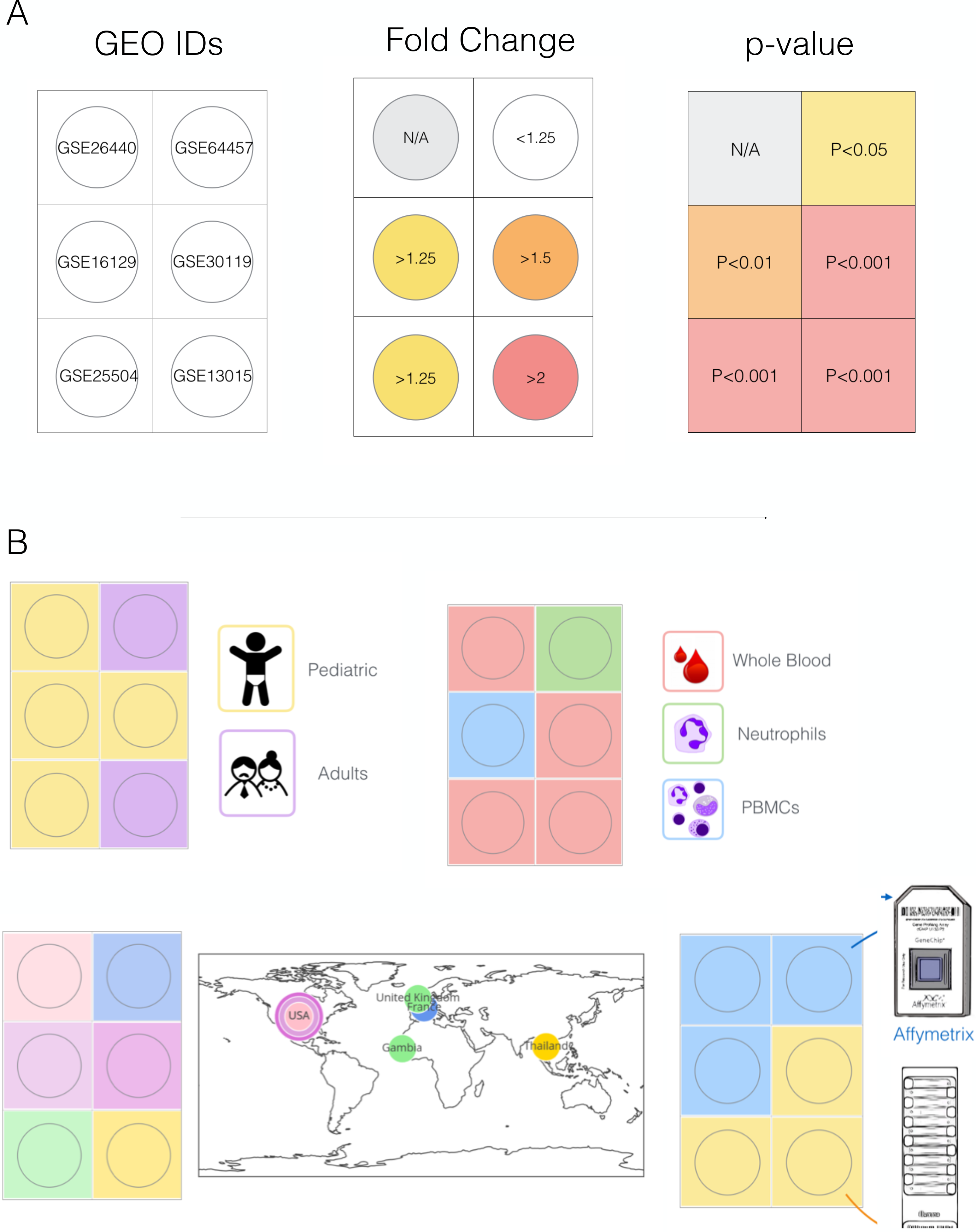
Independent validation of the primary observation. A. NUDT16 fold change and significance levels. Fold change and t-test p-values are shown for datasets which selected for independent validation. B. Characteristics of the validation datasets. Validation datasets were selected from GEO based on their relevance. All involved samples obtained from sepsis patients and non-infected controls. Variation for geographical location, population and biological samples used for profiling are represented graphically.

**Table 1:** List of validation datasets, group information and changes in NUDT16 expression.

| GSE ID | Title | Group A | Group B | Exp A | Exp B | B/A | t-test | F-test |
| --- | --- | --- | --- | --- | --- | --- | --- | --- |
| GSE26440 | Expression data for derivation of septic shock subgroups (130) | Normal Control | Septic shock patient | N/A | N/A | N/A | N/A | N/A |
| GSE30119 | ... whole blood transcriptional response to community-acquired Staphylococcus aureus infection ... (143) | Healthy | S. aureus | 37.5 | 62.4 | 1.7 | <0.001 | <0.001 |
| GSE16129 | Enhanced Monocyte Response & Decreased Central Memory T Cells in Children with Staphylococcus aureus Infection (52) | Ctrl | SA | 716.2 | 960.3 | 1.3 | <0.01 | <0.01 |
| GSE64457 | Marked alterations of neutrophil functions during sepsis-induced immunosuppression (23) | Control | Septic patient | 36.0 | 57.5 | 1.6 | 0.053 | <0.05 |
| GSE13015 | ...A Blood Biomarker Signature for the Diagnosis of Septicemic Melioidosis- (39) | Healthy & T2D | Melioidosis & other sepsis | 39.9 | 84.0 | 2.1 | <0.001 | <0.001 |
| GSE25504 | Whole blood mRNA expression profiling of host molecular networks in neonatal sepsis (63) | Control | Infected | 125.7 | 162.1 | 1.3 | <0.001 | <0.001 |

### RNA decapping activity indirectly links NUDT16 with inflammatory processes

Although the NUDT16 literature does not overlap with the Sepsis literature it may intermediate biological concepts that indirectly links NUDT16 with inflammatory process may be identified.

Biological concepts were first manually extracted from the NUDT16 literature. The NUDT16 query presented earlier was run again, but this time restricting the search to title by using [ti] (instead of [tw] = “text words”). This search returned 8 articles. Keywords were identified in the title of each one of them and categorized (Table 2). Categories included for instance: biomolecules, metals, molecular processes, cell type, tissue, disease. Next the prevalence of these concepts in the overall NUDT16 literature was determined. The NUDT16 PubMed query was run against text words ([tw]) adding in turn each of the keywords. The largest number of PubMed hits recorded was for the keyword “mRNA decapping” with the following query: “*Nudix hydrolase 16” [tw] OR NUDT16 [tw] OR H29K [tw] AND “mRNA decapping”* (8 articles). When searching for “RNA decapping” instead, 4 articles were returned. Combining both “mRNA decapping” AND “RNA decapping” returned 10 articles. A similar strategy was used to identify the next two most prevalent concepts, which were “Inosine Triphosphate” with 4 articles and “Inosine Diphosphate”, with 3 articles.

**Figure 3:**
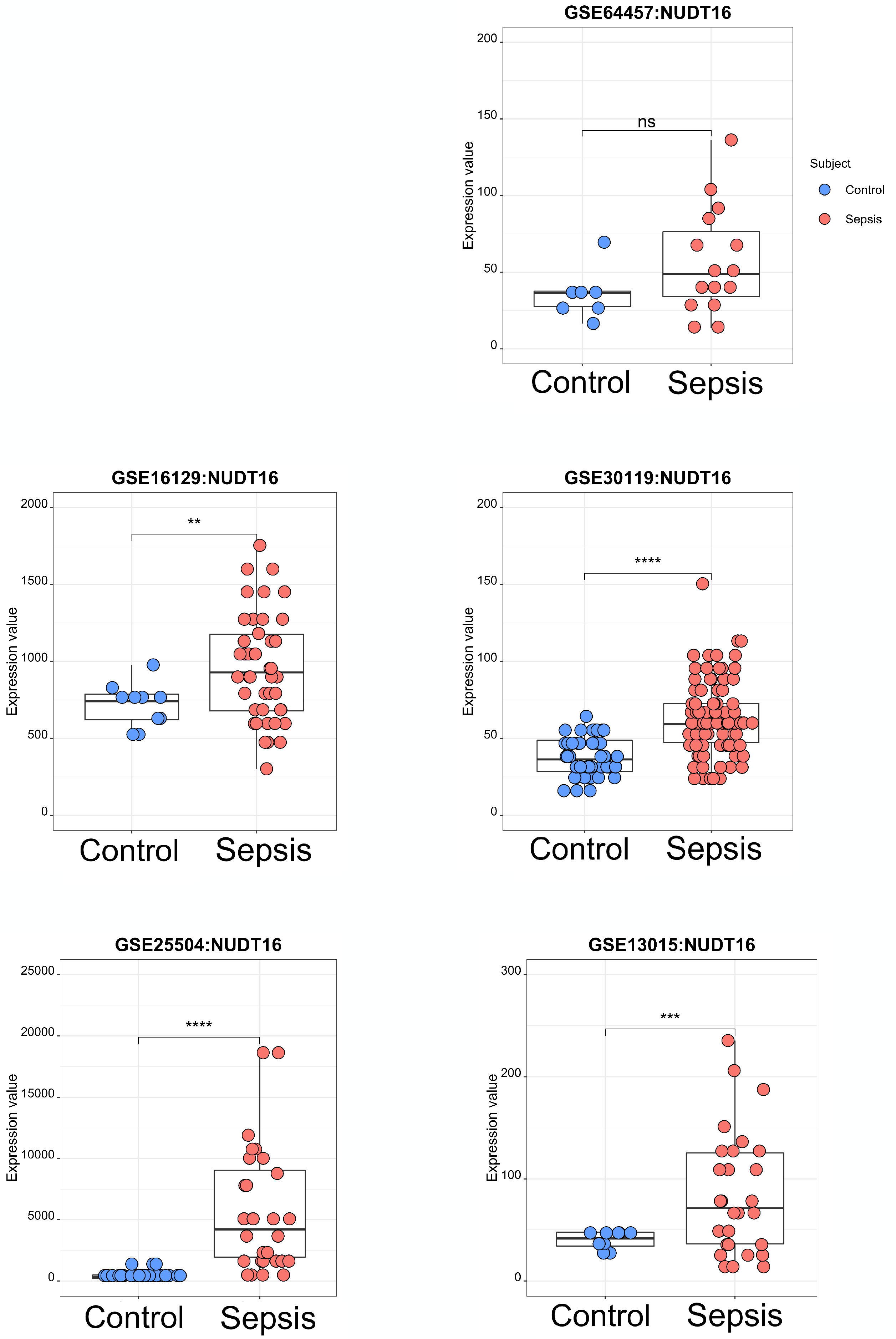
Transcriptional profiles of NUDT16 in datasets selected for validation. Levels of transcript abundance in leukocytes of sepsis and control subjects are shown for six independent validation datasets.

**Table 2:** Concepts extracted from the NUDT16 literature and their prevalence.

| Category | NUDT16-associated Concept | PubMed Articles |
| --- | --- | --- |
| Activity | Hydrolytic activity | 1 |
| Biomolecule | Dinucleotide cap analogs | 1 |
| Biomolecule | Short capped oligonucleotides | 1 |
| Molecular process | mRNA decapping | 8 |
| Biomolecule | C-MYC | 1 |
| Disease | acute lymphoblastic leukemia | 1 |
| Cell type | T-cell | 1 |
| Biomolecule | Inosine monophosphate | 2 |
| Biomolecule | Deoxyinosine phosphatase | 2 |
| Biomolecule | Inosine phosphatase | 1 |
| Biomolecule | ITPA | 4 |
| Molecular process | Chromosome stability | 1 |
| Molecular process | Cell growth | 2 |
| Biomolecule | IDP | 3 |
| Biomolecule | dIDP | 3 |
| Biomolecule | ITP | 4 |
| Biomolecule | dITP | 3 |
| Molecular process | RNA binding | 1 |
| Molecular process | RNA decapping | 4 |
| Metal | Cobalt | 1 |
| Metal | Magnesium | 2 |
| Metal | Manganese | 1 |
|  | "mRNA decapping" OR "RNA decapping" | 10 |
|  | "inosine triphosphate" | 4 |
|  | "inosine diphosphate" | 3 |

The next step consisted in exploring the relevance of these three key NUDT16 biological concepts in the sepsis, inflammation or neutrophil literature. A series of PubMed queries were run, with for example: *(“mRNA decapping” OR “RNA decapping“) AND inflammation* identifying 4 articles linking indirectly NUDT16 with inflammation (**Figure 4**). No articles were found among the sepsis or neutrophil literature that relates to RNA decapping. Similar queries were run against the inflammation, sepsis or neutrophil literature for “inosine triphosphate” (**Figure 4**) or “inosine diphosphate” (**Supplementary Figure 3**). The number of literature articles linking these two NUDT16 biological concepts with inflammation, sepsis or neutrophils ranged between 1 (“Inosine triphosphate” AND sepsis) and 16 (“Inosine diphosphate” AND inflammation).

**Figure 4:**
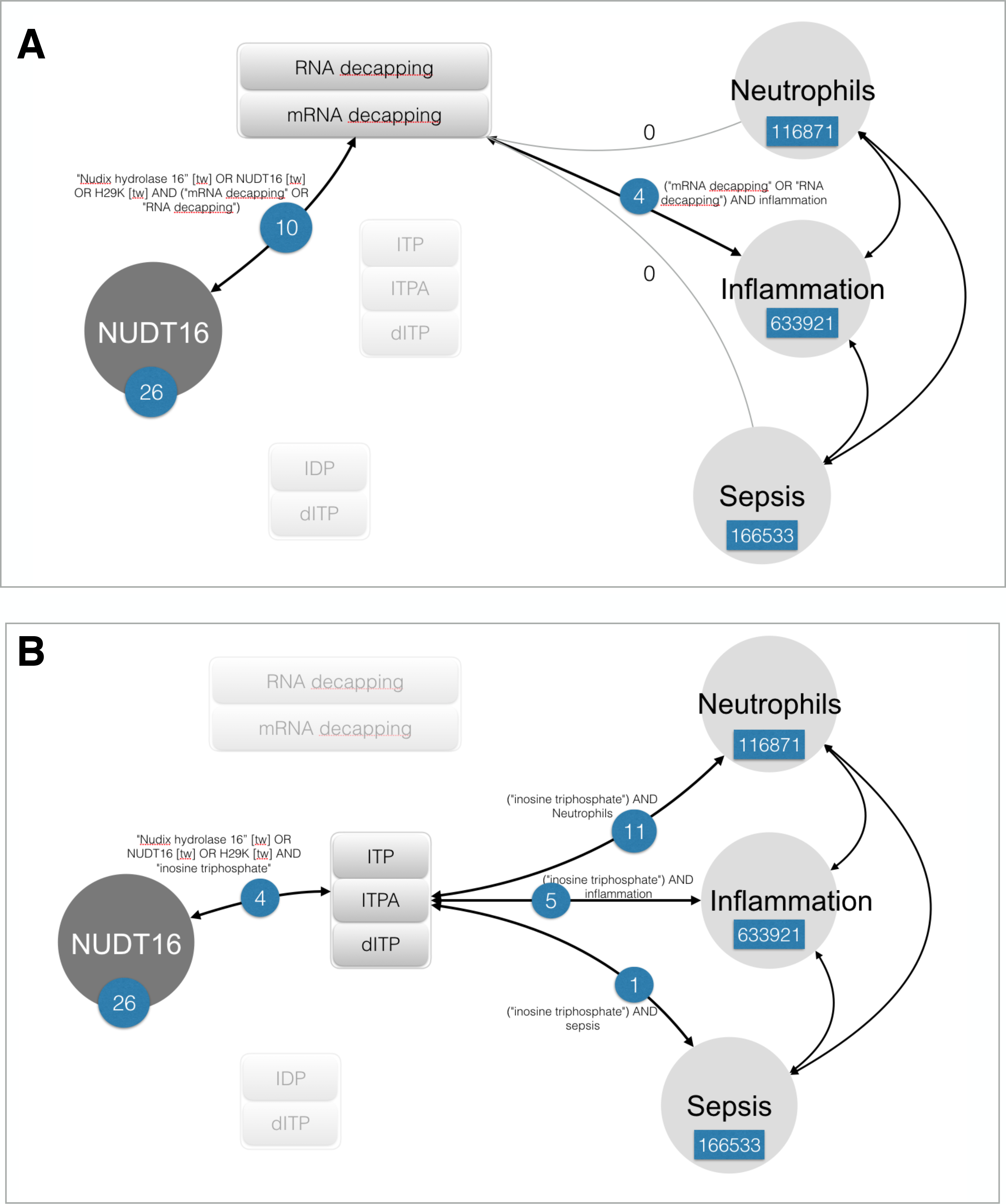
Intermediate concepts linking indirectly NUDT16 to Neutrophils, Inflammation or Sepsis. The most prevalent biological concepts in the NUDT16 literature were used in searches against the neutrophil, inflammation or sepsis literature. The extent of the overlap with de NUDT16 literature is shown here for “RNA decapping”, “Inosine triphosphate”.

Inferences can be made regarding the potential role of NUDT16 in activated neutrophils from the establishment of these indirect links, including: 1) in increasing degradation or turnover of mRNA species via the exonuclease pathway (on the basis of: [2, 11]). 2) in protecting neutrophils from the deleterious accumulation of (deoxy)Inosine diphosphate in the nucleus (on the basis of: [12]).

## DISCUSSION

Inferences made regarding a potential role of NUDT16 in human sepsis would require experimental follow up. For one, to verify whether increase in abundance observed at the transcript level translates in increase in protein abundance. This could be done in an in vitro culture system, exposing neutrophils to serum of sepsis patient as was done in the initial study described in **Figure 1**. Indeed, in this study exposing neutrophils to LPS did not induce NUDT16 expression. It was not the case either in another dataset in which blood was exposed in vitro for 6 hours to a wide range of pathogen- and host-derived immune stimuli ([13]/ GEO accession: GSE30101). Using neutrophils isolated from patients with sepsis would be another possibility but fold changes increase in abundance of NUDT16 transcripts was found to be lower than in the in vitro serum exposure system. Second, ascertaining cellular localization of NUDT16 by confocal microscopy would prove particularly informative since both nuclear and cytoplasmic localization has been reported before, with different functional implications [12, 14]. There are no reports of NUDT16 KO mice having been produced or phenotyped to date. If viable, testing survival of NUDT16 null animals in experimental models of sepsis or inflammation (e.g. endotoxin administration, cecal ligation models) could provide additional insights with regards to its function.

Additional inferences may be made for instance by exploring transcriptional profiles of other members of the Nudix family, including for instance the decapping enzyme DCP2, which activity bears similarities to that of NUDT16 within the exonuclease degradation pathway, while showing a differential spectrum of RNA specificity [3, 15, 16].

## ACKNOWLEDGEMENTS

Sidra Medicine is a member of the Qatar Foundation for Education, Science and Community Development. The work was supported in part via a grant from the Qatar National Research Fund: QNRF NPRP10-0205-170348

## AUTHOR CONTRIBUTION

Conceptualization: MG, DR, DC. Data curation and validation: DR, MT and SB. Visualization: DR, DC. Analysis and interpretation: MG, DR, DC. Writing of the first draft: DC. Funding acquisition: SB, DB, NM, DC. Methodology development: MG, DC, DR. Project administration: DC. Software development: MT, SB. Writing – review & editing: MG, DR, BK, MT, MA, JR, WH, SB, NM, DB, DC. The contributor’s roles listed above follow the Contributor Roles Taxonomy (CRediT) managed by The Consortia Advancing Standards in Research Administration Information (CASRAI) (https://casrai.org/credit/).

---

**Supplementary Figure 1:**
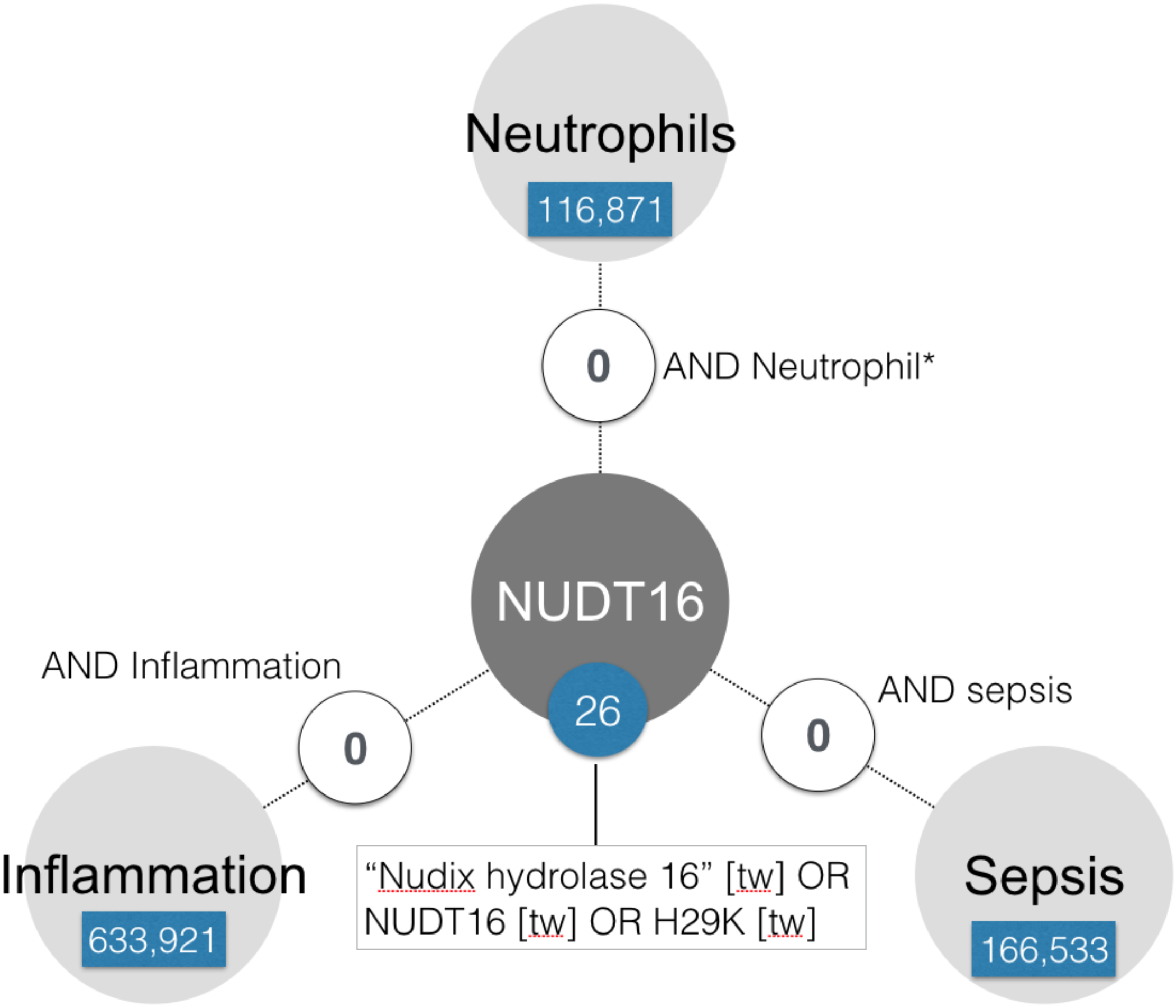
Assessment of gap in knowledge in the biomedical literature. PubMed queries were designed to identify associations between the NUDT16, Sepsis, Neutrophil and Inflammation literature. No overlap was found indicating the probable existence of a gap in biomedical knowledge.

**Supplementary Figure 2:**
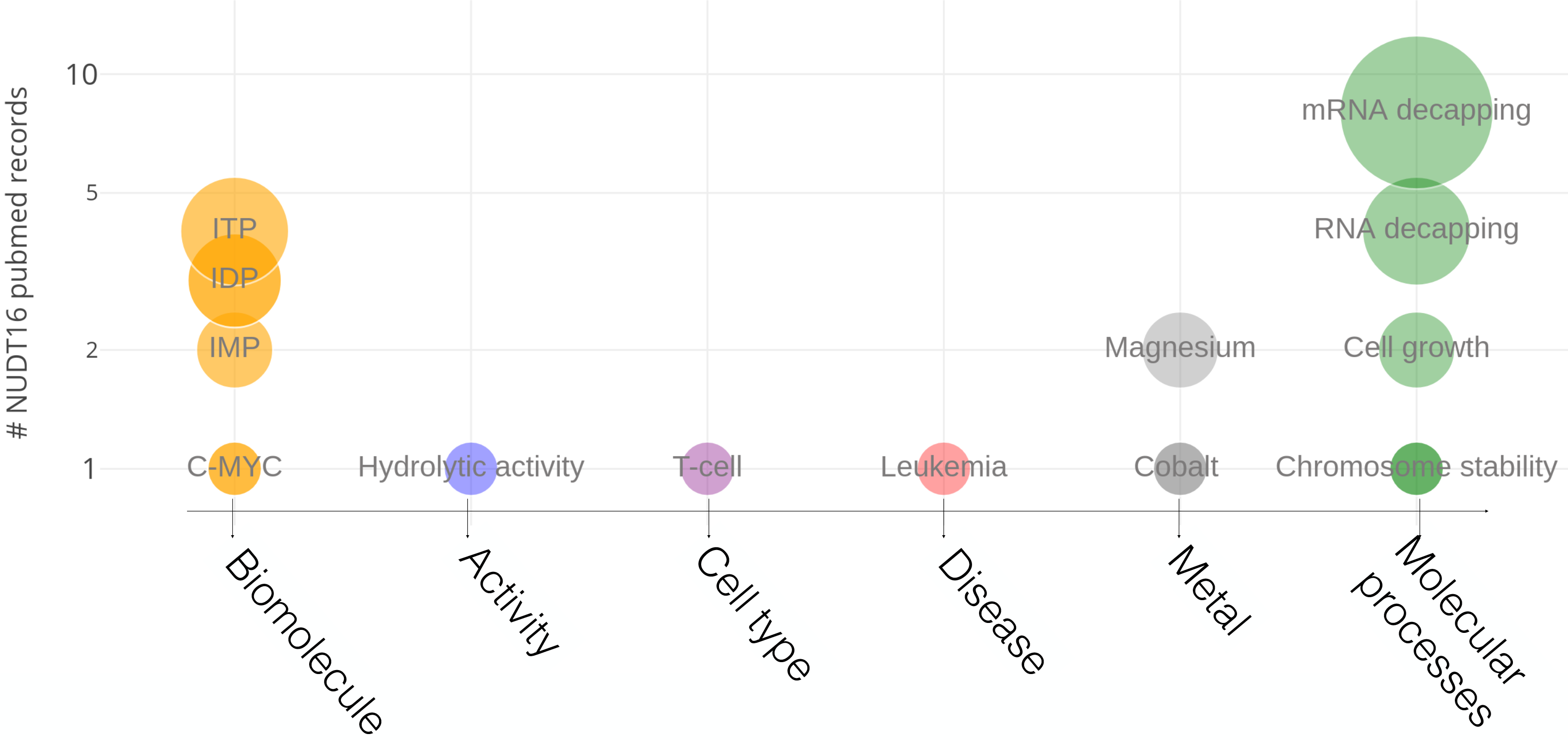
NUDT16 biological concepts. Biological concepts were extracted from the NUDT16 literature which comprises of 26 articles. The frequency of PubMed articles is shown.

**Supplementary Figure 3:**
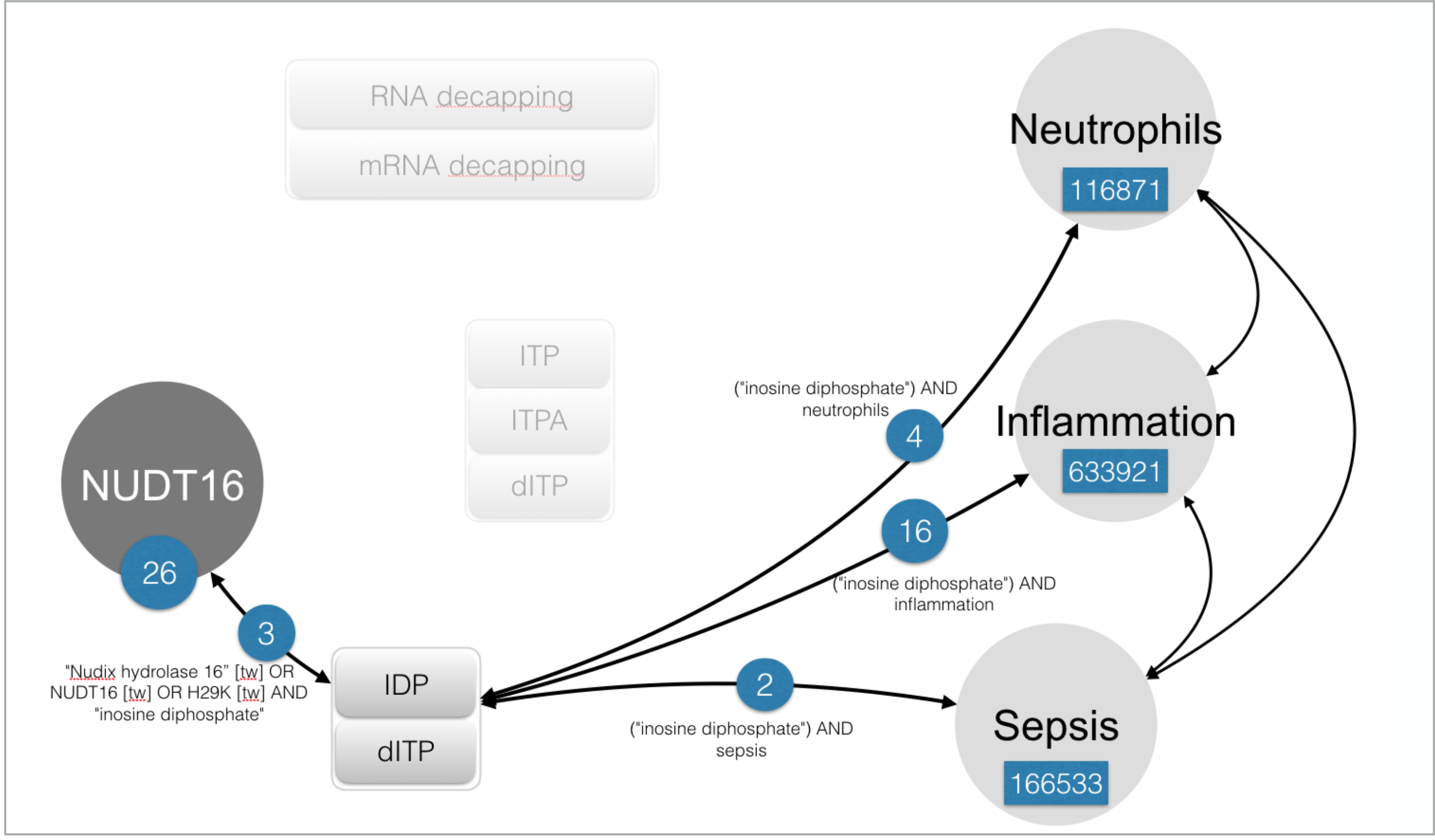
Intermediate concepts linking indirectly NUDT16 to Neutrophils, Inflammation or Sepsis. The most prevalent biological concepts in the NUDT16 literature were used in searches against the neutrophil, inflammation or sepsis literature. The extent of the overlap with de NUDT16 literature is shown here for “Inosine diphosphate”.

